# Mechanistic dissection of antibody inhibition of influenza entry yields unexpected heterogeneity

**DOI:** 10.1101/2022.08.03.502654

**Authors:** Anjali Sengar, Marcos Cervantes, Peter M. Kasson

## Abstract

Neutralizing antibodies against influenza have generally been classified according to their recognition sites, with antibodies against the head domain of hemagglutinin thought to inhibit attachment and antibodies against the stalk region thought to inhibit fusion. Here, we report the development of a microfluidic assay to measure neutralization of viral entry that can clearly differentiate between effects on attachment and fusion. Testing multiple broadly-neutralizing antibodies against the hemagglutinin stalk domain, we obtain a surprising result: some broadly-neutralizing antibodies inhibit fusion only, while others inhibit both fusion and viral attachment. Antibodies binding the globular head domain primarily inhibit attachment but can also reduce the fusogenic capability of viral particles that nonetheless bind receptor. These findings shed light on the unexpectedly heterogeneous mechanisms of antibody neutralization even within similar recognition sites. The assay we have developed also provides a tool to optimize vaccine design by permitting assessment of the elicited antibody response with greater mechanistic resolution.

## Introduction

Influenza continues to pose a threat to public health, causing 1 billion infections, 3 to 5 million cases of severe illness, and 290,000 to 650,000 deaths throughout the world every year (1, 2). Antigenic evolution of the virus makes preventing influenza challenging: vaccines require seasonal redesign based on recent circulating strains and have highly variable efficacy. Therefore, longstanding research efforts have aimed to develop more universal influenza vaccines that elicit broadly effective neutralizing antibody responses to prevent seasonal and pandemic influenza. To accomplish this, however, better means of measuring neutralization mechanisms are needed.

Hemagglutinin (HA), a major influenza surface glycoprotein, is a major antigenic target for both vaccine-elicited and infection-elicited immunity. HA is synthesized as a single polypeptide chain (HA0) that assembles into a trimer (3). Host tissue proteases subsequently cleave HA0 into HA1 and HA2 (4). HA1 subunit solely forms the hemagglutinin “head” and mediates viral attachment to sialic acid cell receptors on target cells, while HA2 forms the majority of the hemagglutinin “stem” and undergoes low-pH-induced conformational changes to drive fusion of the viral membrane with the host membrane (5, 6). Most neutralizing antibodies target epitopes at or near the receptor-binding site on the HA head and generate strain-specific responses (7). A small fraction of these head-binding antibodies, such as CH65 (7), CR8033 (8), and C05 (9–11), can neutralize a broad spectrum of influenza subtypes. However, the majority of broadly neutralizing antibodies target the HA stem region. These include CR6261 (12, 13), F10 (14), CR8020 (13), FI6 (15), MEDI8852 (16) and CR9114 (8).

The current understanding of antibody neutralization mechanisms is largely guided by structure-function relationships and numerous crystallographic or electron cryomicroscopy structures of antibody-HA complexes (8, 13, 15). Most high-resolution Fab-HA structural studies have concluded that direct neutralization occurs by blockade of receptor binding or by restricting the conformational changes required for membrane fusion. Traditional cell-based neutralization assays such as microneutralization, inhibition of viral plaque formation, and hemagglutination inhibition measure the convolved effects of multiple mechanisms of action, including antibody-mediated aggregation of viral particles, inhibition of receptor-binding, inhibition of membrane fusion, and inhibition of progeny virus release. It has thus been challenging to correlate high-resolution structural information with precise functional roles, although some important efforts have been made in this regard (17–19).

Here, we have developed a way to further dissect mechanisms of antibody neutralization against influenza by differentiating receptor binding, fusion by bound viruses, and fusion of total viruses. To accomplish this, we use single-virus fluorescence microscopy (20) and two assays: one where influenza virus binds to sialic-acid receptors, and a second where virus is captured using synthetic DNA-lipid conjugates and made accessible for fusion in a receptor-independent fashion (Fig. 1A) (21). Influenza virions are fluorescently labeled, incubated with different IgG antibodies, and then measured for their ability to bind and fuse to synthetic liposomes displaying either model glycosphingolipid receptors or DNA-lipids. These data are analyzed to yield four parameters: inhibition of virus binding to receptors, inhibition of fusion by receptor-bound virus, inhibition of fusion by total virus independent of receptor-binding capacity, and antibody binding to the whole virion in the same conditions.

**Figure 1:**
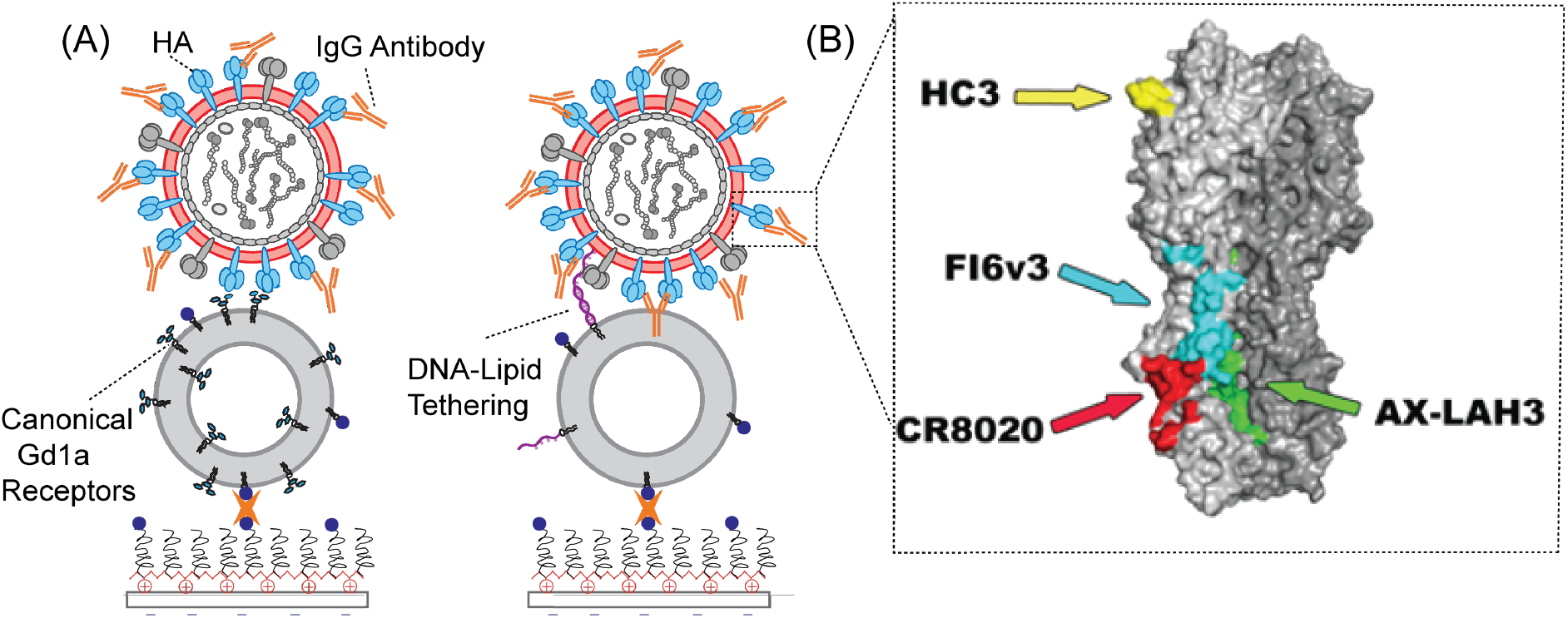
(A) Schematic of influenza binding and fusion assays. Fluorescently labeled viral particles bind target liposomes via either sialic-acid-containing GD1a receptors (left) and or hybridization of complementary DNA strands tethered to the viral membrane and the target vesicle membrane (right). Virus is pre-incubated with antibodies against hemagglutinin. (B) A surface rendering of H3 hemagglutinin in its pre-fusion state (PDB 1HGF) is colored to show the binding footprint of antibodies studied: HC3 (yellow), CR8020 (red), AX-LAH3 (green), and FI6v3 (cyan).

These measurements have the benefit of revealing unanticipated second mechanisms for neutralization by some antibodies. For instance, we found that some stem-binding antibodies inhibit receptor binding as potently as they inhibit fusion, while others primarily affect fusion. Conversely, antibodies against the receptor-binding site can also slow fusion kinetics at higher antibody coverage. This may result from the different abilities of bivalent IgG antibodies to bind to sterically available HA epitopes on the surface and cross-link inter and intra -HA trimers, inhibiting or delaying productive fusion. These findings may help explain other reports that concluded not all hemagglutinin molecules must be bound in order to prevent an influenza virus from undergoing membrane fusion. The resulting insights into antibody mechanisms of action together with structural understanding may aid the development of better vaccines and antivirals.

## Results

Here we examined the interaction of four IgG antibodies targeting different sites on hemagglutinin (Fig. 1B): HC3 (22, 23), AxLaH3 (24), CR8020 (8), and FI6v3 (15). HC3 targets an unusual protruding loop in the receptor-binding domain on HA1 residues (140-146). AX-LAH3 was prepared against the long alpha region (LAH) HA2 residues (74-128) recognizing the stalk domain from H3 strains. CR8020 targets residues on the HA2 stem near the fusion peptide and membrane-proximal G helix (15-16, 18-19, 25, 30, 32-36, 38, 146, 150), and a single HA1 residue (325). FI6v3 targets HA1 residues (8, 28, 29, 30, 287, 289-290, 316) and HA2 residues (18-21, 38-39, 41-43, 45-48, 49, 52-53, 56-57) in fusion subunit on the HA stalk. FI6v3 was reported to approach the HA stem with very different angles and orientations, recognizing helix A and spanning to the fusion peptide of the next monomer.

First, we assayed the antibody neutralization of infectivity using MDCK2 cell monolayers as described previously (25). HC3 exhibited the most potent neutralization activity (EC_50_ ∼0.3 μg/mL) compared to the stem binding antibodies FI6v3 (EC_50_ ∼4.8 μg/mL) and CR8020 (EC_50_ ∼43 μg/mL). AX-LAH3 did not achieve 50% inhibitory activity in concentrations as high as 50 μg/mL (Fig 2).

**Figure 2:**
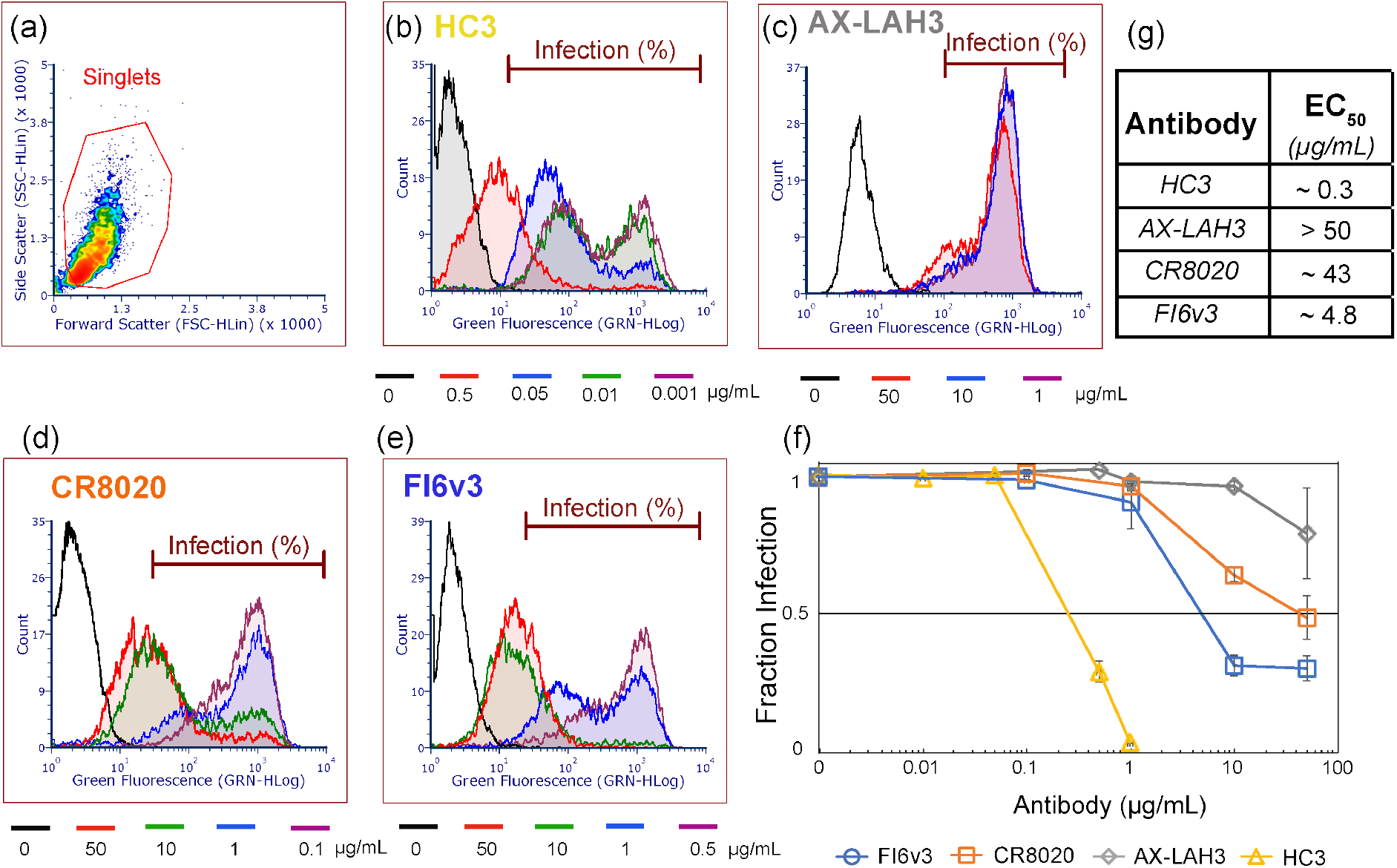
Antibody inhibition of MDCK cell infection by influenza virus. Viral samples were pre-incubated with the designated antibody and then added to MDCK2 monolayers, fixed and stained 6 hours post-infection, and analyzed for hemagglutinin expression via flow cytometry. As shown in panel (a), all samples were gated to select singlet cell populations. Histograms of hemagglutinin expression (green fluorescence from Alexa 488-labeled antibody) are plotted for HC3 (b), AX-LAH3 (c), CR8020 (d), and FI6v3 (e). Histograms for multiple antibody concentrations are overlaid in different colors. The fraction of hemagglutinin-expressing and thus virus-infected cells is plotted in (f) as a function of antibody concentration, normalized to no antibody control. Error bars indicate standard deviations. Estimated EC_50_ values extracted from that plot are listed in (g).

We also probed the binding of monoclonal antibodies to X-31 (H3N2 A/Aichi/68) influenza virions using immunofluorescence. Antibody-virus complexes were allowed to bind to the GD1a receptor vesicles inside the microfluidic flow cell and stained using secondary antibodies. The number of colocalized Texas-red labeled virions with secondary antibodies was calculated to obtain the binding affinity values (EC_50_) for different antibodies. EC_50_ values were measured as 1.1 μg/mL for HC3, 0.6 μg/mL for CR8020, 0.73 μg/mL for FI6v3, and 27.5 μg/mL for AX-LAH3 (Fig S1). Both head binding (HC3), and stem binding (CR8020, FI6v3) antibodies exhibited comparably high-affinity binding, whereas AX-LAH3 bound weakly. It is worth noting that this assay only measures virions that can bind receptors, so in the case of HC3 the EC_50_ may be shifted slightly due to the high-potency inhibition of viral attachment.

Single-virus assays of viral binding and fusion provide a mechanistic complement to functional assays of cell infection. To measure antibody inhibition of viral binding and fusion, we pre-incubated X-31 influenza virus with each of the antibodies and then added the resulting solution to a microfluidic flow cell coated with liposomes containing either a) GD1a sphingolipid model receptors or b) DNA-lipids (in this case with complementary DNA-lipids previously added to the virus) to measure binding. Measuring binding in these two modes differentiates (a) blockade of sialic-acid receptor binding by hemagglutinin versus (b) steric blockade of viral attachment by other means. Subsequent to this, fusion was triggered via a pH drop, and both the fraction of bound viruses undergoing lipid mixing and the kinetics of lipid mixing were assessed. In GD1a-binding mode, these measurements report on fusion inhibition of viral particles that are able to bind receptor, whereas in DNA-binding mode, these measurements report on fusion inhibition of all available virus, irrespective of the sialic-acid binding activity.

We first present the results on inhibitory EC_50_ values grouped by antibody. The HC3 monoclonal antibody efficiently inhibited viral binding to GdD1a receptors with an EC_50_ of 6.5 µg/mL (Fig. 3A). Virions that succeeded in binding GD1a were further inhibited for fusion with an EC_50_ of 7.55 µg/mL, suggesting that virions that had incomplete sialic-acid-binding blockade were also impaired for fusion. HC3 did not measurably inhibit viral binding in DNA-binding mode, showing no blanket steric blockade of the viral surface. Most interestingly, it also did not inhibit viral fusion in DNA-binding mode at 10 µg/mL (Fig. 3B). Together, these results suggest that HC3 primarily inhibits receptor binding by influenza as previously suggested but that it can also inhibit fusion by the subset of viral particles that have bound HC3 but are still capable of binding, while most of the binding-incapable particles can still fuse efficiently if bound in a receptor-independent manner. We did not observe any significant delay in mean hemifusion time as a function of HC3 binding (Fig. 3C, Fig. 3D).

**Figure 3:**
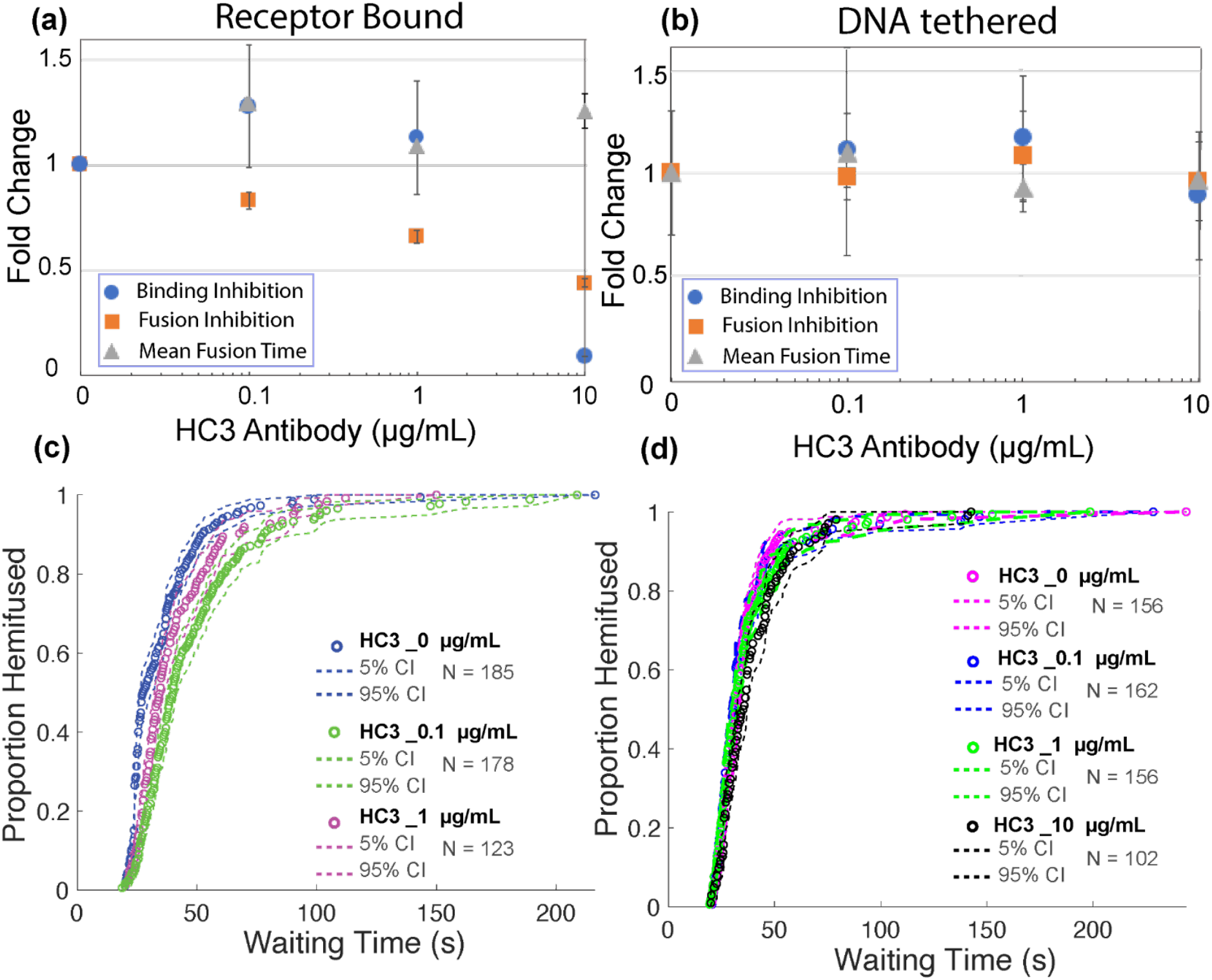
Effect of HC3 antibody on influenza virus binding inhibition, fusion inhibition, and mean hemifusion time. Values denote fold-change relative to no antibody samples with increasing concentration of antibody. Error bars indicate standard deviations. Inhibition is plotted for (a) GD1a receptor binding mode, (b) DNA tethering mode, (c) lipid mixing kinetics of virus bound to GD1a receptors, and (d) lipid mixing kinetics virus bound via DNA-lipid tethering. Cumulative distribution functions (CDFs) for the waiting time from pH drop to single-virus lipid mixing are plotted together with 95% confidence intervals from bootstrap resampling.

The first stalk-binding antibody tested, CR8020, also inhibited influenza binding to liposomes with an EC_50_ of 36 µg/mL. There was also some reduction of binding by DNA-tethered virus, although no clear dose-response relationship was evident. Inhibition of fusion by receptor-bound virus was observed with an EC_50_ of 34.4 µg/mL, while fusion by DNA-tethered virus was inhibited with an EC_50_ of 7.4 µg/mL (Fig. 4A, Fig. 4B). These data demonstrate a previously underappreciated dual mode of action for CR8020 in inhibiting both binding and fusion. Incubation with CR8020 also induced a delay in mean hemifusion times: a 1.86-fold delay was measured at 50 μg/mL CR8020 in Gd1a-binding mode and 1.5-fold at 10 µg/mL CR8020 in DNA-tethering mode (Fig. 4C, Fig. 4D).

**Figure 4:**
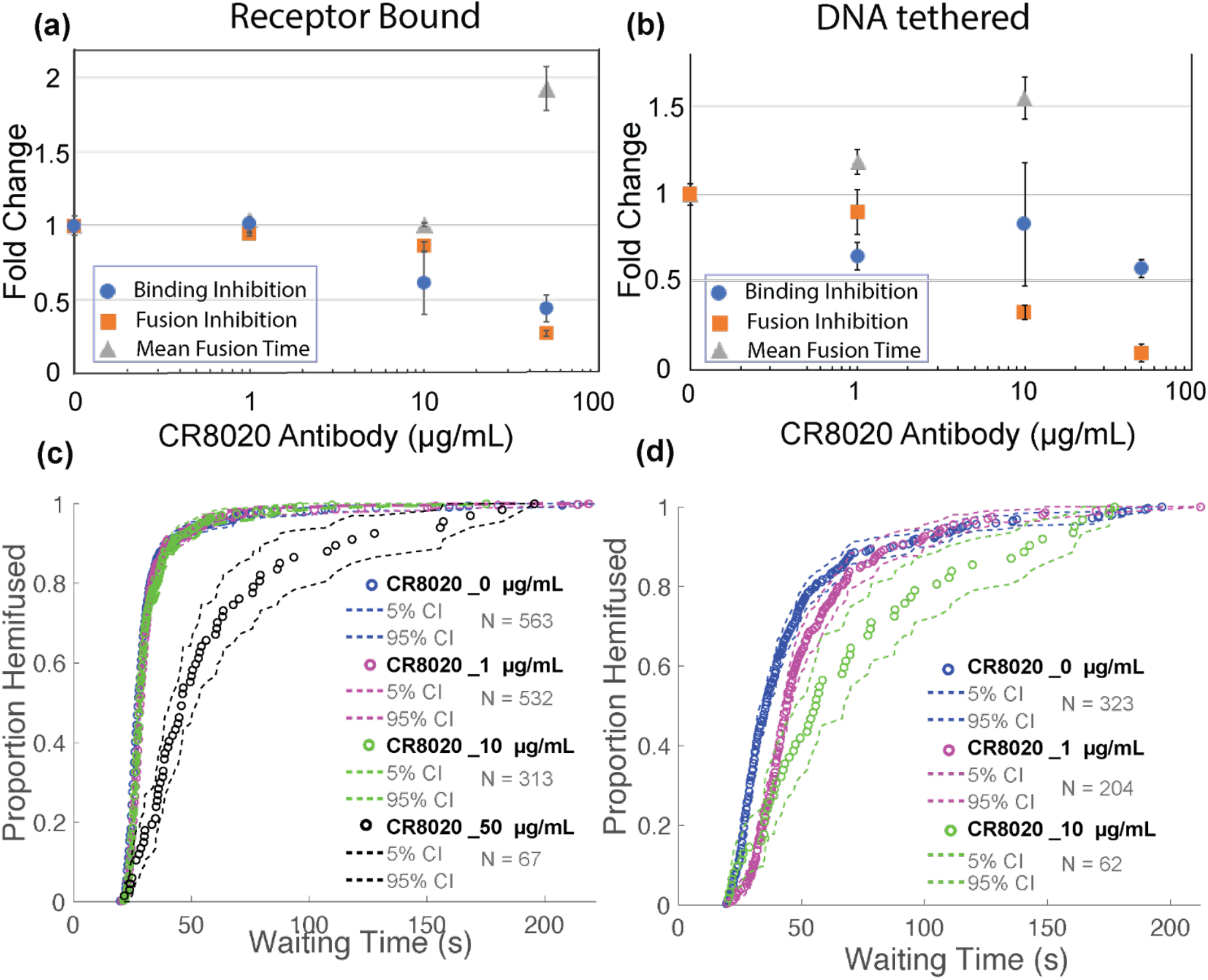
Effect of CR8020 antibody on influenza virus binding inhibition, fusion inhibition, and mean hemifusion time. Values denote fold-change with increasing antibody concentration, and error bars indicate standard deviations. Inhibition is plotted for (a) receptor binding mode, (b) DNA tethering mode. (c) lipid mixing kinetics of virus bound to GD1a receptors, and (d) lipid mixing kinetics virus bound via DNA-lipid tethering. CDFs for the waiting time from pH drop to single-virus lipid mixing are plotted together with 95% confidence intervals from bootstrap resampling.

In contrast to CR8020, the FI6v3 antibody did not measurably inhibit viral binding either to GD1a receptors or in DNA-tethering mode. FI6v3 did show potent fusion inhibition, with EC_50_ values of 4.3 µg/mL for GD1a-bound virus and 12.5 µg/mL for DNA-tethered virus (Fig. 5A, Fig. 5B). A delay in mean hemifusion times was also observed: 2.2-fold at 50 µg/mL FI6v3 in Gd1a binding mode and 1.8-fold change at 10 µg/mL FI6v3 in DNA tethering mode. These are consistent with a purely fusion-inhibitory effect (Fig. 5C, Fig. 5D).

**Figure 5:**
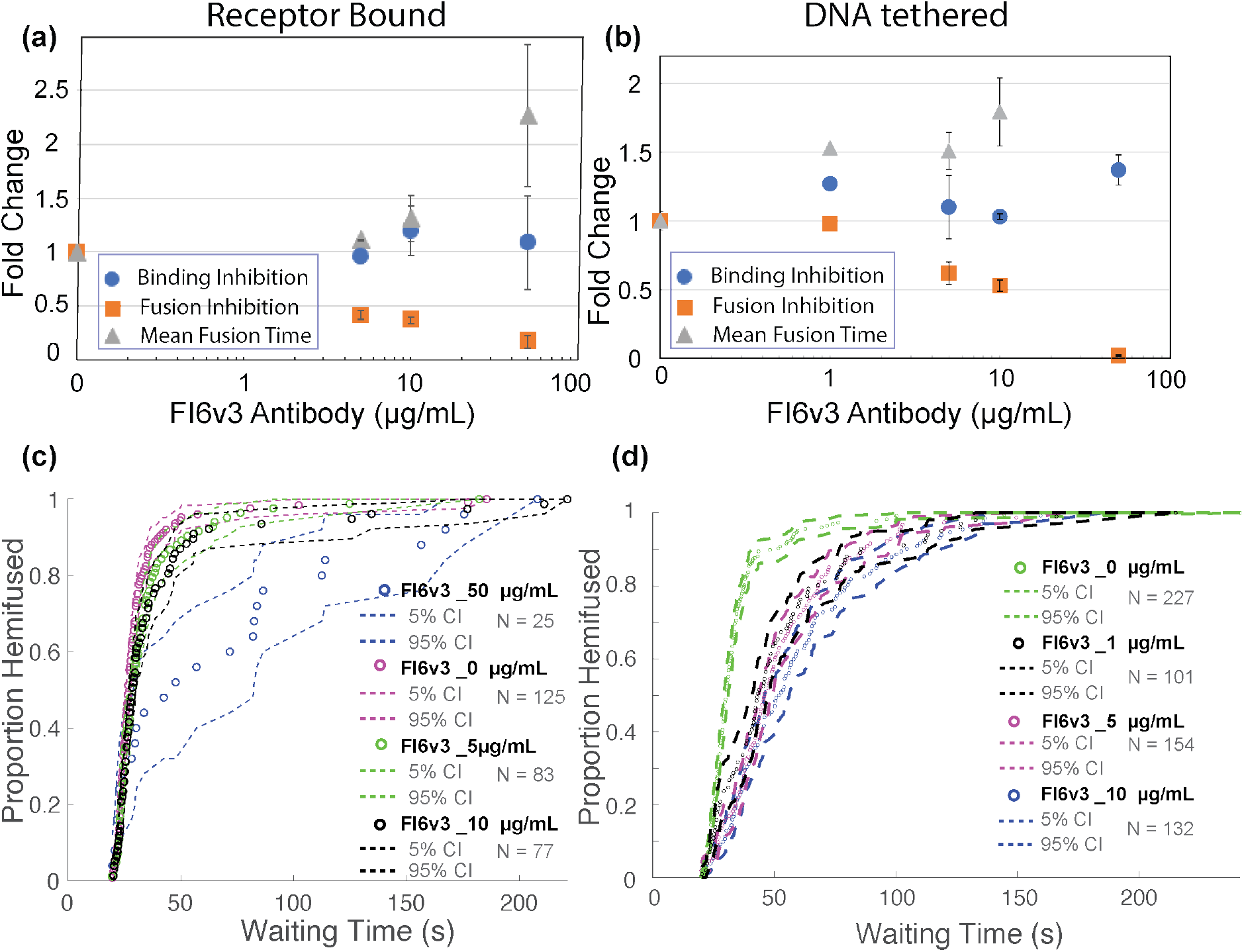
Effect of FI6v3 antibody on influenza virus binding inhibition, fusion inhibition, and mean hemifusion time. Values denote fold change with increasing antibody concentration, and error bars indicate standard deviations. Inhibition is plotted for (a) receptor binding mode, (b) DNA tethering mode. (c) lipid mixing kinetics of virus bound to GD1a receptors, and (d) lipid mixing kinetics virus bound via DNA-lipid tethering. CDFs for the waiting time from pH drop to single-virus lipid mixing are plotted together with 95% confidence intervals from bootstrap resampling.

The AX-LAH3 antibody displayed much weaker binding (EC_50_ 27.5 µg/mL) to viral particles than other antibodies tested. Consistent with this, single virus studies showed that AX-LAH3 did not substantially inhibit binding either to GD1a or with DNA-lipids (EC_50_ > 50 µg/mL, Fig. S2). Similarly, AX-LAH3 did not detectably inhibit fusion at concentrations as high as 50 μg/mL or slow lipid mixing times (Fig. 6). Given that other antibodies showed inhibitory EC_50_ values at greater than five times their binding EC_50_, these findings are not surprising, but they do serve as a control to exclude non-specific binding or fusion inhibition by immunoglobulins in our assay.

**Figure 6:**
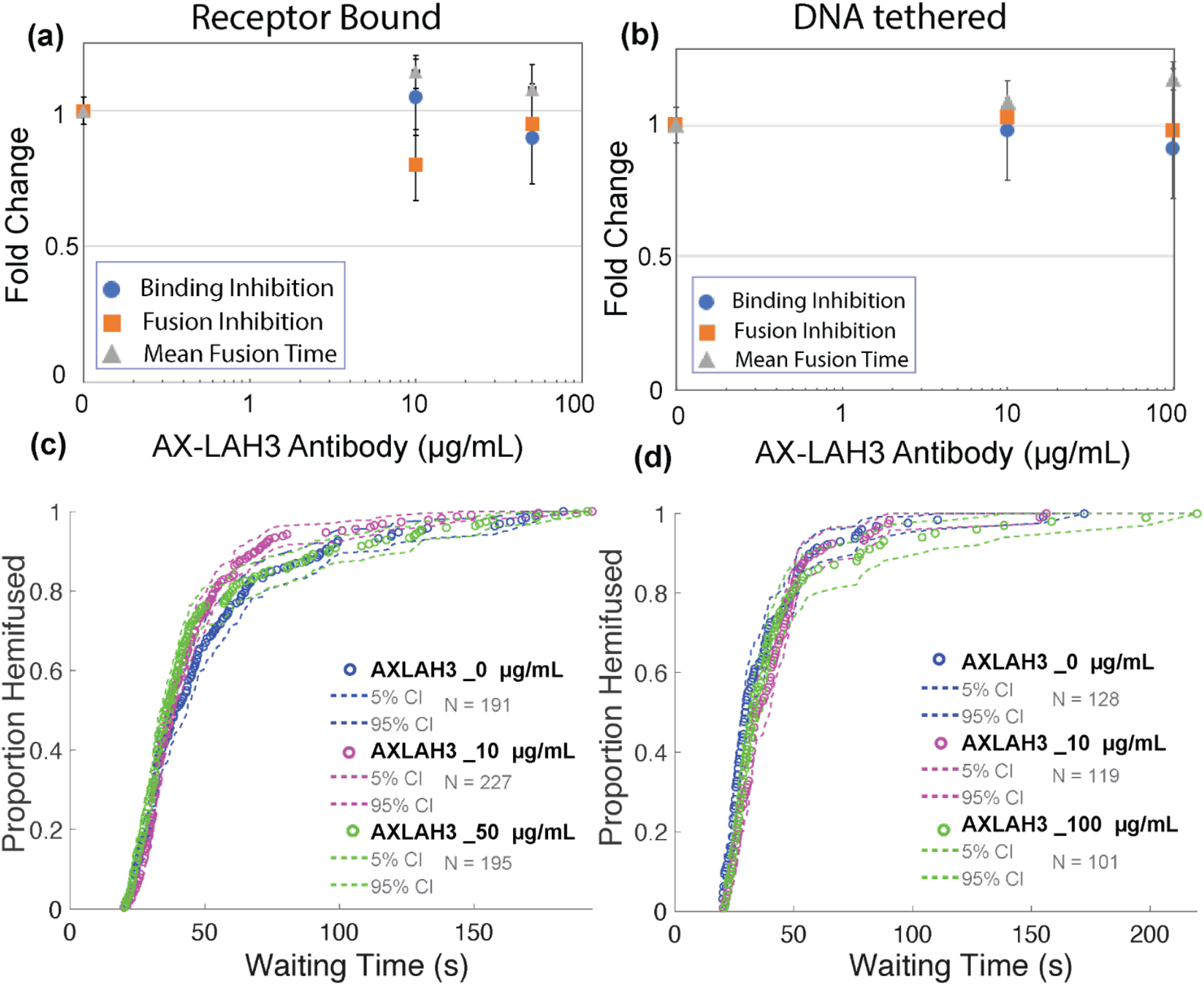
Effect of AX-LAH3 antibody on influenza virus binding inhibition, fusion inhibition, and mean hemifusion time. Values denote fold change with increasing antibody concentration, and error bars indicate standard deviations. Inhibition is plotted for (a) receptor binding mode, (b) DNA tethering mode, (c) lipid mixing kinetics of virus bound to GD1a receptors, and (d) lipid mixing kinetics virus bound via DNA-lipid tethering. CDFs for the waiting time from pH drop to single-virus lipid mixing are plotted together with 95% confidence intervals from bootstrap resampling.

## Discussion

We describe a method to differentiate antibody inhibition of viral attachment versus inhibition of viral fusion. The particular advantage of our approach is that it can also test whether viral particles inhibited for attachment are fusion-competent, leading to more precise mechanistic insight into antibody activity. We used HC3 as a model attachment-blocking antibody since it has an epitope that overlaps with the receptor-binding site on HA1 (23). We observed that increased HC3 concentration greatly decreased the number of viruses that bind target membranes displaying sialic acid receptors. This is consistent with the known mechanism for head-binding antibodies to inhibit virus attachment (26). Surprisingly, we also observed that HC3 could inhibit fusion by the subset of HC3-bound viral particles that could nonetheless bind target membranes. Based on this, we conclude that the number of available hemagglutinin molecules required for viral attachment is likely lower than the number required for membrane fusion, at least in the case of HC3 inhibition. An additional factor could be the fraction of hemagglutinin cleaved into HA1/HA2 complexes and thus available for fusion, as discussed by Ivanovic and colleagues (27). Analogous effects have been previously observed for HC19 (28), another head-binding antibody, but in that case, the effect was attributed to the ability of bivalent IgG to cross-link intraparticle or interparticle HA trimers (19). HC19 was demonstrated to form clusters by crosslinking individual virus particles in electron microscopy, which provided an explanation for the fusion inhibition observed by bulk fluorescence assay. Our single-virus fusion inhibition data, on the other hand, is selective for single viruses eliminating the possibility of interparticle cross-linking of HA, leaving only the possibility of crosslinking on the same particle. Fusion inhibition may be caused by head-binding antibodies interference with the rearrangement of the HA1. Both the “uncaging” (29–32) and the “fusion peptide release” (31, 33, 34) models ultimately require HA1 reorganization prior to membrane fusion. Such a mechanism of fusion inhibition may be extended to other head-binding antibodies; results may depend on the antibody binding location and the angle of approach, which could affect the ability of bivalent IgG to link HAs on a trimer or from adjacent trimers. Interestingly, viral morphology was recently found to have an effect on antibody inhibitory concentrations for membrane fusion (28), suggesting additional dimensions to this comparison.

The two stem-binding, broadly neutralizing antibodies CR8020 and FI6v3 showed a large difference in their potencies to neutralize infection in MDCK2 cells: FI6v3 was approximately nine times more potent than CR8020 (Fig 2). In our single virus fusion assay, FI6v3 was found to be 7-8 times more effective than CR8020 at inhibiting fusion of GD1a-receptor-bound virus, while both antibodies were equally effective at inhibiting fusion of virus captured via receptor-independent DNA tethering (Fig S3). This provides an informative contrast in their neutralization mechanism. Since virions with high occupancy of bound CR8020 appear unable to bind GD1a-decorated membranes, the fusion inhibition assay using receptor-bound-virus should select for virions with low antibody coverage and thus display a higher EC_50_ than DNA-tethering capture, which will assay virions containing the full range of antibody coverage. However, their binding EC_50_ values to HA displayed on virions were similar: FI6v3 was slightly more potent than CR8020 (Fig S1). This dual action of CR8020 in inhibiting attachment to GD1a-decorated membranes with almost equal potency to fusion inhibition is surprising. This unanticipated dual effect of CR8020 in inhibiting binding as well as fusion was also noticed by Otterstorm et al. at high antibody concentrations (>20 µg/mL/>150 nM), but the magnitude of binding inhibition was not quantified (17). Other reported effects of CR8020 on the efficiency of fusion inhibition and delay in hemifusion time in the same study were found to be in the same range as ours.

Some insights about the differences in CR8020 and FI6v3 binding to HA are provided by prior research on antibody structures and competitive inhibition (15, 35–37). Slight differences in antibody-bound structures are consistent with a different mechanism of action, although we would have predicted these to be too subtle to have functional effects. The FI6v3 epitope sits slightly higher up (central-stalk epitope) on the HA stem compared to the CR8020 epitope, which is located at the base of HA (proximal epitope), close to the viral membrane. Structural studies also reveal a markedly different angle of approach for bound antibodies between CR8020 and FI6v3. Despite these tantalizing glimpses, the mechanisms by which CR8020 affects receptor binding but not FI6v3 remain unclear and require further investigation.

In summary, we show how antibodies against influenza virus can have subtle, unanticipated functional effects. Differentiating inhibition of viral attachment, inhibition of fusion by receptor-bound virions, and inhibition of fusion by all virions in a sample can yield additional insight into antibody mechanism and highlights how broad structure-function generalizations can miss subtler structural effects with important functional consequences, particularly in the case of viral fusion glycoproteins that adopt multiple conformations over the course of attachment and entry.

## Materials and Methods

### Reagents

Palmitoyl phosphatidylcholine (POPC), dioleoyl phosphatidylethanolamine (DOPE), and Biotin-PE were purchased from Avanti Polar Lipids (Alabaster, AL). Cholesterol and disialoganglioside (GD1a) from bovine brain (Cer-Glc-Gal(NeuAc)-GalNAc-Gal-NeuAc) were purchased from Sigma-Aldrich (St. Louis, MO). Texas Red-1,2-dihexadecanoyl-sn-glycero-3-phosphoethanolamine (TR-DHPE), Oregon Green-1,2-dihexadecanoyl-sn-glycero-3-phosphoethanolamine (OG-DHPE) and Neutravidin were purchased from ThermoFisher Scientific (Waltham, MA). Polydimethylsiloxane (PDMS) was obtained from Ellsworth Adhesives (Hayward, CA). Poly(L-lysine)-graftpoly(ethylene glycol) (PLL-g-PEG) and Poly(L-lysine)-graft-poly-(ethylene glycol) biotin (PLL-g-PEG biotin) were purchased from SuSoS (Dübendorf, Switzerland). Influenza A virus (strain X-31, A/Aichi/68, H3N2) was purchased from Charles River Laboratories (Wilmington, MA), and all experiments using virus were performed under BSL-2 safety conditions with institutionally approved protocols. Primary antibodies are as follows: AX-LAH3 (monoclonal antibody to influenza A H3 hemagglutinin (HA) long alpha helix 3 in the stalk domain) was obtained from BEI resources NR-50513, CR8020 (recombinant human anti-H3 HA antibody) was purchased from Creative Biolabs (Shirley, New York), FI6v3 (recombinant monoclonal IgG against HA stalk domains) was a gift from Jesse Bloom, and HC3 (monoclonal IgG against H3 HA receptor-binding domain) was a gift from J. Skehel. Secondary antibodies are as follows: Alexa Fluor 647 donkey anti-mouse IgG (H+L) cat # A31571, Alexa Fluor 647 goat anti-human IgG (H+L) cat # A21445, Alexa Fluor 488 mouse anti -human IgG1 cat # A10631, Alexa Fluor 488 goat anti-mouse IgG (H+L)/IgM (L) cat # A28175, all purchased from Invitrogen (Waltham, Massachusetts).

### Buffers

The following buffers were used: Reaction buffer: 10 mM NaH_2_PO_4_, 90 mM sodium citrate, and 150 mM NaCl (pH 7.4); fusion buffer: 10 mM NaH_2_PO_4_, 90 mM sodium citrate, and 150 mM NaCl (pH 5.0); Hepes buffer (HB): 20 mM HEPES and 150 mM NaCl (pH 7.2).

### DNA Sequence and tethering

DNA sequences with 5’ lipids were custom ordered from IDT DNA. DNA Sequence ‘C’ used to tether liposomes is /5DPPEK/TA GTA TTC AAC ATT TCC GTG TCGA and complementary DNA sequence ‘D’ incorporated in viral particles is /5DPPEK/TT TTT TTT TTT TTT TTT TTTTTT TTC GAC ACG GAA ATG TTG AAT ACTA with T24 linker as previously used (21). Lyophilised DNA sequences were diluted to the 100 μM concentration in a buffer with 2 mM HEPES, and 15 mM NaCl at pH 7.2 (10x dilution of Hepes Buffer), stored at -20 °C.

### Vesicle preparation and DNA incorporation

Large unilamellar vesicles (LUV) were made following established protocols (38). Lipids (67% POPC, 20% DOPE, 10% cholesterol, 1% biotin-DPPE and 2% GD1a, total 140 nM lipids) were mixed in chloroform. For liposomes without Gd1a, 69% POPC was used instead, and for a subset of samples used to calibrate the timescale of pH drop, 0.5% pH-sensitive fluorescent dye OG-DHPE was added, leaving 66.5% POPC. Solvent was evaporated by a gentle stream of nitrogen to form a thin film, which was stored under house vacuum overnight. The dried lipid film was then hydrated in a reaction buffer, incubated for 30 min, and vortexed for 5-10 min to obtain a lipid suspension at a concentration of 0.56 mM lipid. This was subjected to 5 freeze-thaw cycles to obtain unilamellar vesicles, 21 extrusion passes through polycarbonate membrane filters (Avestin) with pore diameter of 100 nm. Vesicles without Gd1a were incubated with lipid-DNA ‘sequence C’ at 0.05 mol % of the total lipid concentration overnight at 4 °C to allow DNA tether incorporation. For DNA tether experiments vesicles were incubated with lipid-DNA Sequence-C. For 100 µL of vesicles, 1 µL of 6 µM ‘sequence C’ DNA-lipid was mixed and incubated overnight at 4 °C. Vesicles were used within one week of preparation.

### Texas Red labeling of Influenza and DNA incorporation

For single virus fusion experiments, X-31 influenza virus (H3N2 A/Aichi/68 HA and NA) was labeled with the fluorescent membrane label Texas Red-DHPE following a protocol previously used for influenza and Zika viruses (21),(39),(40). Briefly, Texas Red solution (0.74 mg/mL) in ethanol was diluted in the HB buffer at a ratio of 1:40. 100 µL of influenza solution (viral protein concentration from Charles River ∼ 0.2 mg/mL) was mixed in 400 µL of diluted Texas Red-DHPE suspension in HB buffer and incubated at room temperature in the dark for 2 h on a rocker. The resulting virus-dye mixture was diluted with 15 mL of HB buffer, and divided into 10 centrifuge tubes each 1.5 mL. Each aliquot was pelleted by spinning at 20,000 x g for 60 min at 4 °C, pellets from all tubes were resuspended and collected in a total of 100 µL HB buffer. Labeled virus was then aliquoted and stored at -80 °C until use. For DNA tether experiments labeled viral particles were incubated with lipid-DNA Sequence-D. For 15 µL of Texas-Red labeled virus, 1 µL of 1 µM ‘sequence D’ DNA-lipid was mixed and incubated overnight at 4 °C. This labeled virus-DNA mixture was diluted 10x for fusion experiments and maintained no more than one week at 4 °C.

### Microfluidic Flow Cell Preparation

To prepare microfluidic flow cells, we used the protocol described before (21). Briefly, glass coverslips (24 × 40 mm, No 1.5, VWR International) were cleaned using a detergent:water mixture in a ratio of 1:7 (7x detergent MP Biomedicals) with continuous heating and stirring for 30 min, until the solution turned clear. This was followed by rinsing extensively in DI water, baking for 4 hours at 400 °C in a kiln, and overnight slow cooling. Coverslips were then rinsed with ultrapure water and sonicated in ethanol for 10 min. Cleaned coverslips were again rinsed and sonicated in water and ethanol and dried in a 100 °C incubator. Cooled coverslips were then stored in a 50 mL falcon tube and wrapped with parafilm until use.

Microfluidic flow-cell molds were prepared using tape-based soft lithography (41). by affixing Kapton polyimide tape (Ted Pella) on a microscopic slide to make channels of dimensions 1 mm x 13 mm x 70 µm. Polydimethylsiloxane (PDMS; Sylgard 184) and catalyst were mixed in a 1:10 ratio and poured into a mold. After degassing under a house vacuum, the PDMS was cured at 60 °C for 4 hours, and flow cells were cut out. Channel inlet and outlet holes were created using a 2 mm biopsy punch.

Glass coverslips were plasma cleaned for 5 mins (Harrick Plasma) before bonding. PDMS flow cells and cleaned glass coverslips were plasma bonded together after plasma activation for 1 min. Immediately after, PDMS flow cell channels were coated with 95% PLL-PEG: 5% PLL-PEG-biotin. After 30 min incubation, channels were washed with 1 mL of ultrapure water and 1 mL of Hepes buffer (20 mM Hepes, 150 mM NaCl, pH 7.2). The biotin-PEG layer was functionalized by incubating for 15 min with a 0.2 mg/mL solution of neutravidin (Thermo Scientific). Channels were washed with 1 mL of HB to remove excess neutravidin. 10 µL of biotin-functionalized liposomes were added to the flow cell channel and incubated for 2 h at RT or overnight at 4 °C for biotin neutravidin binding; these two methods yielded equivalent liposome coverage. Each flow cell channel was washed with 1 mL of HB buffer to remove unbound vesicles.

### Neutralizing Antibodies

For each experiment, 10 µL of labeled influenza virus was mixed with the antibodies in the designated concentration and incubated for 45-60 min at room temperature, then added to the flow cell channel.

### Lipid-Mixing Assay

Each microfluidic flow cell (coated with liposomes) was washed and equilibrated using Reaction Buffer. To initiate the experiment, 10 µL of antibody-treated influenza virus solution was added to the channel and incubated for 30 min for GD1a-decorated liposomes and 1-2 h for DNA-tethered liposomes, reflecting the faster on-rate of GD1a binding than DNA tethering. Channels were washed with 1 mL of Reaction buffer to wash unbound particles. Afterwards, microfluidic flow cells were secured on the microscope stage to stabilize the buffer exchange pump system. Low pH buffer (Reaction Buffer at pH 5.0) was added, and a stream of 1000 images was collected at a frame rate of 0.25 sec/frame. The time between pH exchange and fluorescence dequenching was analyzed as the waiting time to lipid mixing using code described below. Buffer exchange time to pH drop was calibrated using pH-sensitive OG-DHPE vesicles in separate experiments as done previously (21).

### Neutralization assay in MDCK2 cells

MDCK2 cells were cultured at 37 °C, 5% CO_2_ in high glucose Dulbecco’s modified eagle medium (Gibco, cat # 11965092) supplemented with 10% fetal bovine serum (Gibco, cat # 11965092), 1% L-glutamine (Gibco, cat # 25030081), 1% sodium pyruvate (Gibco, cat #11360070). 125,000 cells were seeded into each well of 24-well plates 18-24 h before the experiment to yield ∼90% confluence. For infection, X-31 influenza virus was diluted in Opti-MEM (Gibco, cat # 11058021) at a ratio of 1:10000 to add 250 µL into each well (25). For experiments testing antibody neutralization, the virus was incubated with different antibodies at indicated concentrations for 45 min prior to the experiment at room temperature. After adding virus-antibody mixture, plates were centrifuged at 250 x g for 30 min at 4 °C to permit viral binding and then incubated at 37 °C, 5% CO2 for 6 h to allow for infection and HA protein expression at the cell surface (25)(42). Cells were de-adhered with Accumax (Innovative Cell Technologies, cat # AM105), pelleted at 800 x g, 4 °C, washed twice, and blocked with 3% BSA in phosphate-buffered saline with calcium and magnesium (DPBS, Gibco, cat # 14040133) for 15 min on ice. To assay for surface-expressed HA, cells were incubated with the HC3 antibody (diluted 1:500 in DPBS, 3% BSA) for 45 min on ice, washed and stained with an Alexa 488 anti-mouse secondary antibody (1:1000 diluted in DPBS, 3% BSA) for 30 min on ice. Cells were then fixed with 4% paraformaldehyde for 15 min at room temperature and measured using flow cytometry (Guava easyCyte, EMD Millipore).

### Microscopy

Video micrographs were acquired using a Zeiss Axio Observer inverted microscope using a 100X oil immersion objective. A Spectra-X LED Light Engine (Lumencor) was used as an excitation light source with excitation/emission filter sets as follows: Cy5 (Chroma 49009 ET-Cy5), Texas Red (Chroma 49008), DiO/FITC (Chroma 49011). Images were recorded with an Andor Zyla sCMOS camera (Andor Technologies) using 2×2 binning, controlled using Micromanager (43) software.

### Immunofluorescence assay

Influenza particles, pre-incubated with different antibodies were allowed to bind to Gd1a receptors on the target lipid vesicles inside a microfluidic chamber. Flow cell channels were then washed to get rid of unbound particles followed by blocking with 3% BSA in reaction buffer for 60 min. Secondary antibodies [Alexa 647-goat anti-mouse or Alexa 647 Goat anti-Human IgG (H+L) Cross-Adsorbed (Invitrogen)] at 1:1000 dilution were then incubated for 60 minutes at RT, washed to get rid of excess secondary antibody. Texas Red (viral particles) and Alexa 647 (stained antibodies) images were collected sequentially at the same FOV.

### Data Analysis

Image and time-series analysis was performed using code written in Matlab and available on Github at https://github.com/kassonlab/micrograph-spot-analysis as previously reported (21).

Fusion efficiency is determined in single viral fusion experiments as (# of fused particles/# bound particles ×100) in a field of view. The fold change for fusion and binding inhibition was calculated with respect to no antibody control from the same set of experiments. Two or more experiments were used to calculate the median and standard deviation. Half-maximal inhibitory concentration (EC_50_) values were estimated by simple interpolation.

Flow cytometry data from 5000 single cells were analyzed using FCS Express 7 Research software. Histograms from uninfected wells were gated to represent ∼ 0% infection and wells infected with no antibody were averaged to represent ∼ 100% infection. Duplicate wells were used for calculating mean and standard deviation. Fraction inhibition was calculated as (% infection with no antibody – % infection of the sample) / (%infection with no antibody – % infection from uninfected wells). The half-maximal EC_50_ values were estimated using simple interpolation; where good neutralization was observed, 4-parameter fits yielded similar values.

## Acknowledgements

The authors thank R. Rawle, A. Villamil Giraldo, and G. Morbioli for helpful discussions. This work was supported by grants from the Commonwealth Health Research Board (207-01-18) and the UVA Global Infectious Diseases Institute.

