## Supplementary Information for "Mechanistic dissection of antibody inhibition of influenza entry yields unexpected heterogeneity"

**Supplementary Information for**
**Mechanistic dissection of antibody inhibition of influenza entry yields unexpected heterogeneity.**

Anjali Sengar[1], Marcos Cervantes[1], and Peter M. Kasson[1,2]
1. Departments of Molecular Physiology and Biomedical Engineering, University of Virginia, Charlottesville Virginia 22908 USA.
2. Department of Cell and Molecular Biology, Uppsala University, Uppsala 75105 Sweden.

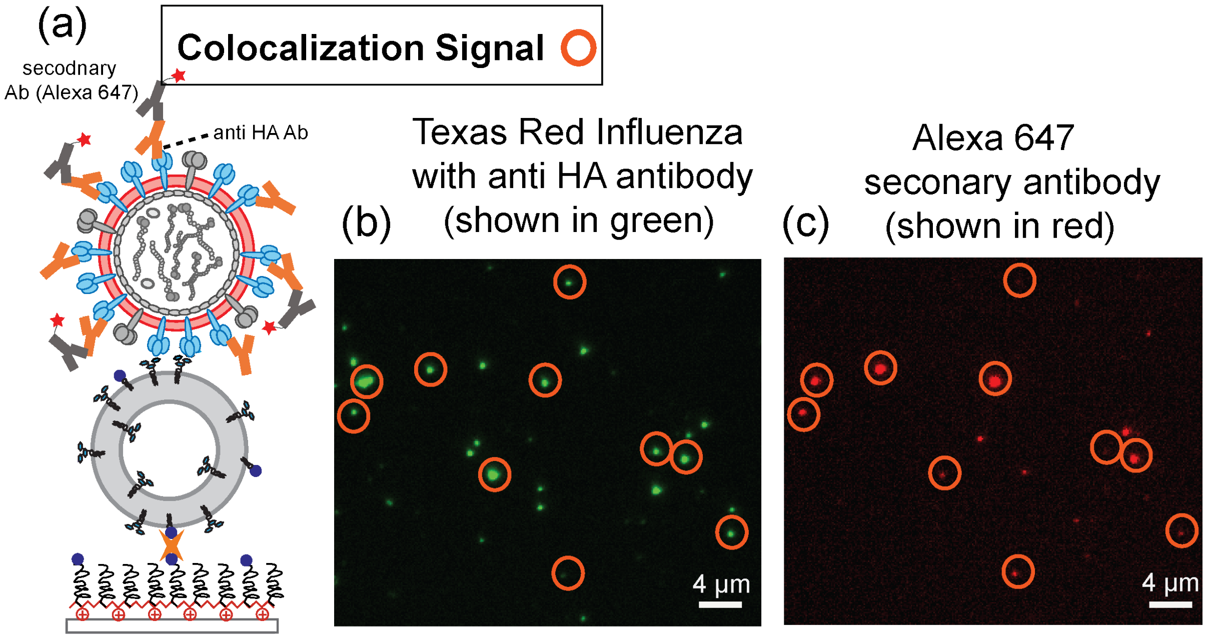


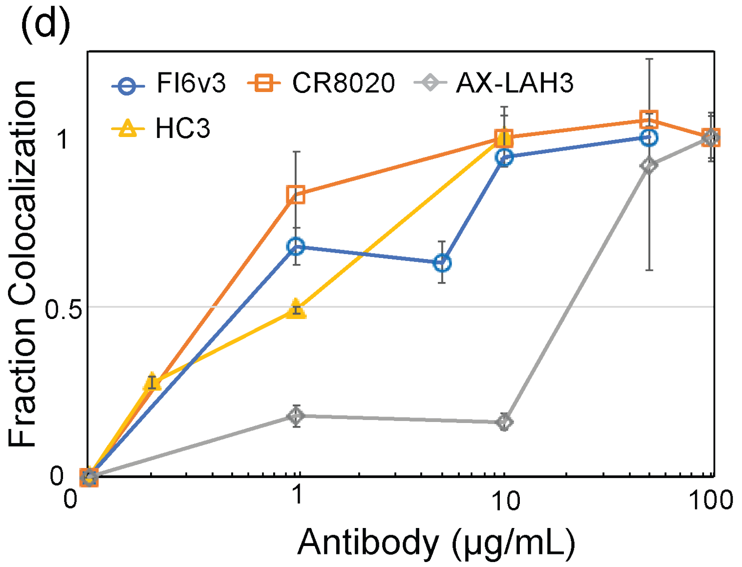


(e)

| Antibody | $EC_{50}$ (μg/mL) |
|---|---|
| *HC3* | ~ 1.1 |
| *AX-LAH3* | ~ 27.5 |
| *CR8020* | ~ 0.6 |
| *FI6v3* | ~ 0.73 |

**Fig S1: Antibody binding to individual viral particles measured via immunofluorescence**. To estimate the binding affinity of different antibodies to influenza viral particles, an immunofluorescence assay was performed, schematized in (a). Texas-Red-labeled viral particles were incubated with different anti-HA antibodies, bound to liposomes displaying GD1a receptor, and stained with Alexa-647 secondary antibody. Sequential images were taken in the Texas Red channel (b) to measure virus and the Alexa-647 channel (c) to measure antibody. Orange circles over particles show co-localized antibody and virus staining. Plotted in (d) is the fraction of labeled virions that also stain positive for antibody with increasing concentration of each antibody with error bars indicating standard deviation. Shown in (e) are estimated binding $EC_{50}$ values for each antibody. Because the virus was bound after antibody incubation, the $EC_{50}$ values represent the convolution of antibody binding and inhibition of receptor binding. For all antibodies except HC3, the inhibition of receptor binding is much less potent and thus in practice not a factor.

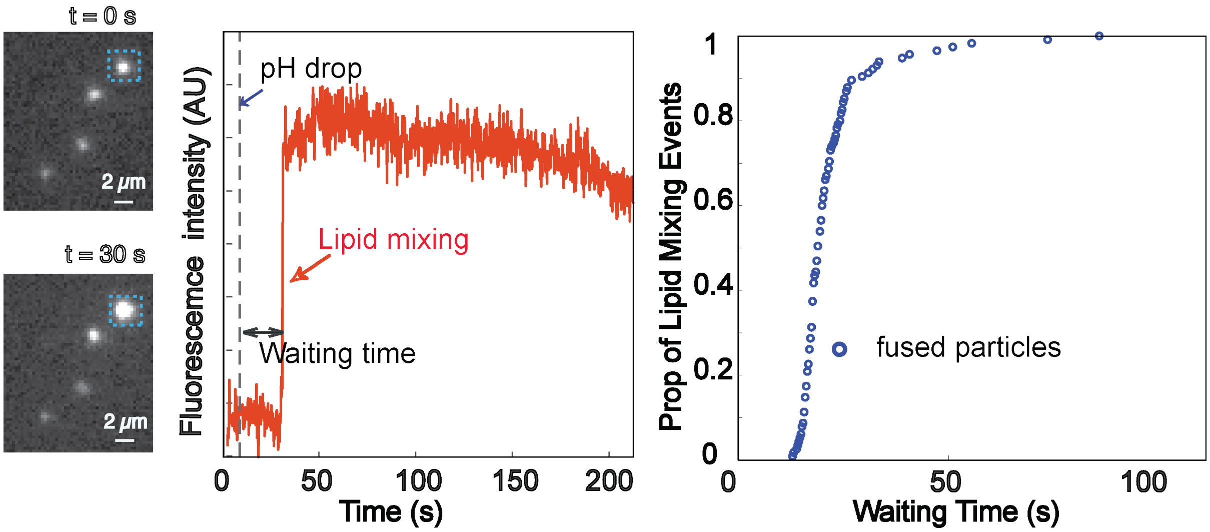


**Fig S2: Sample micrographs, fluorescence intensity trace, and CDF for influenza virus fusion.** Displayed here are a representative micrograph of single-virus spots at the time of pH drop and 30 s after, a fluorescence trace of the particle outlined in blue showing a Texas-Red dequenching event representing lipid mixing, and an assembled CDF of many such particles showing the kinetics of lipid mixing.

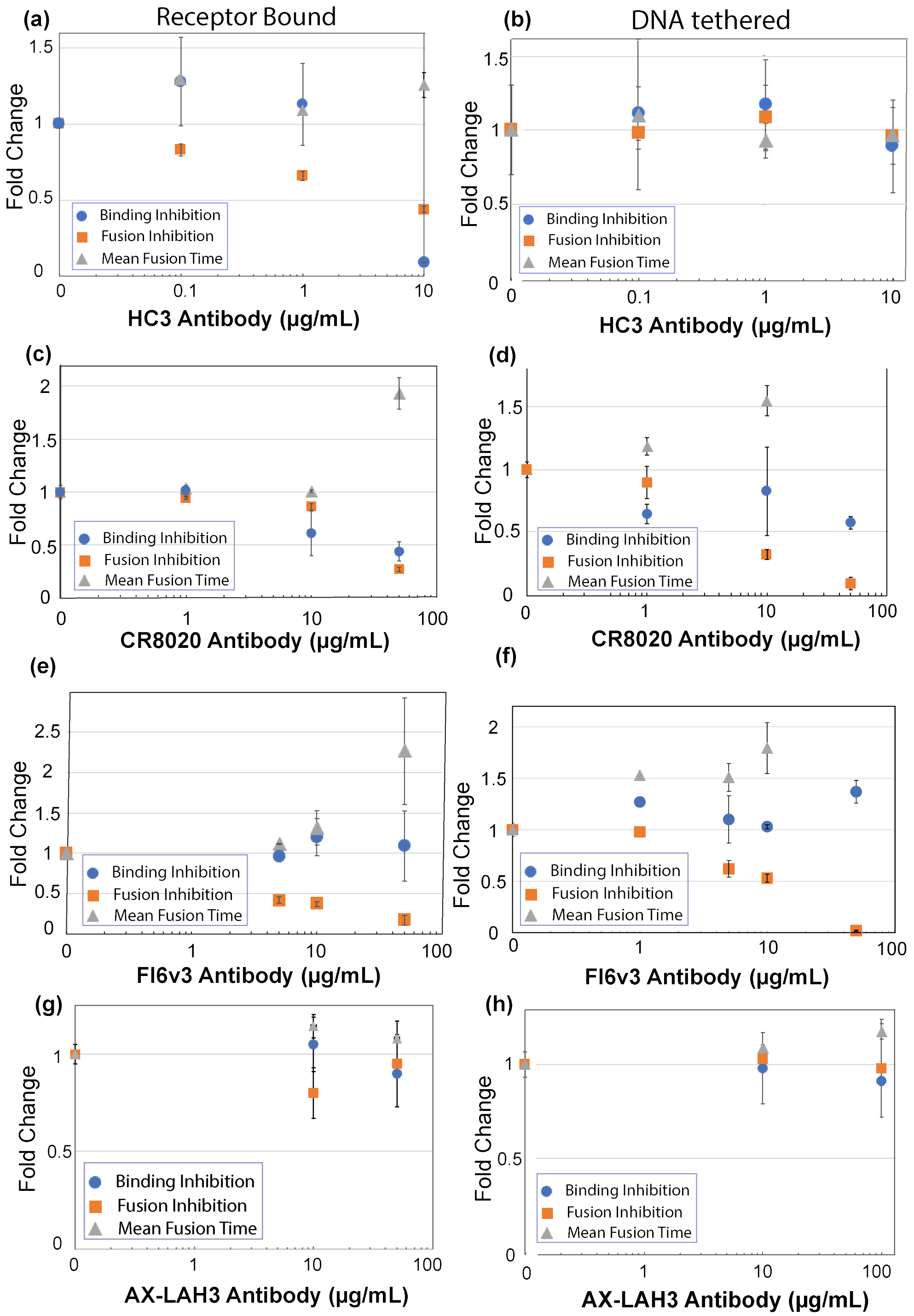


**Fig S3: Effect of different antibodies on influenza virus binding inhibition, fusion inhibition, and mean hemifusion time**. Values denote fold change with increasing antibody concentration, and error bars indicate standard deviations. Inhibition is plotted for (left column) receptor binding mode, (right column) DNA tethering mode. Plots show results for antibodies HC3 (a) and (b), CR8020 (c) and (d), AX-LAH3 (e) and (f) and CR8020 (g) and (h).
